# Endolysosomal acidification amplifies JNK and IMD/NF-κB signaling and exacerbates oxidative injury in the *Drosophila* intestine

**DOI:** 10.64898/2026.07.28.740369

**Authors:** Doug Terry, Liping Luo, Josh Lee, Brian Robinson

## Abstract

Vacuolar ATPases (V-ATPases) are highly conserved multi-subunit proton pumps that drive the acidification of intracellular vesicles, especially endosomes and lysosomes. Beyond enabling cargo degradation, endolysosomal acidification shapes signal transduction, acting both positively through receptor processing in endosomes and negatively through lysosomal turnover of pathway mediators. Whether endolysosomal acidification shapes the intestinal epithelial response to oxidative injury remains unclear. Here we use *Drosophila* to define the role of V-ATPases in intestinal injury driven by excessive oxidative stress. Consistent with lysosomal biogenesis programs induced by oxidant in other systems, we find that oxidant exposure increases V-ATPase subunit expression and abundance of acidified vesicles in the midgut. Yet this response is not protective. RNAi depletion of multiple V-ATPase subunits suppresses lethality induced by either hydrogen peroxide or paraquat exposure. These effects map to mature absorptive enterocytes rather than to intestinal progenitors. Enterocyte-specific depletion of *Vha44* (subunit C of the V1 domain) preserves intestinal barrier function and suppresses the cell death, JNK pathway activation, and IMD/NF-κB reporter activity induced by oxidant exposure. Overexpression of the MAP3K *Tak1* sensitizes animals to oxidative stress and drives JNK and IMD signaling, both of which require *Vha44*. Together, these findings identify endolysosomal acidification as an amplifier of pro-apoptotic JNK and IMD/NF-κB signaling, such that attenuating V-ATPase levels in mature enterocytes protects the intestinal epithelium from excessive oxidant burden.

## Introduction

Oxidative stress has long been associated with inflammation, cancer, and aging [1–3]. Low levels of reactive oxygen species (ROS) production (e.g., H_2_O_2_) have established roles in signaling pathways that influence cell death or survival, differentiation, and migration [2, 4]; while high levels of ROS can cause collateral damage to host tissue driving apoptosis and inflammation [5]. Excessive ROS predominantly comes from the host innate immune response and dysfunctional mitochondria [6–9]. Within the intestine, chronic oxidative stress is believed to play a key role in inflammatory bowel disease (IBD), including Crohn’s disease (CD) and ulcerative colitis (UC) [10, 11]. This chronic oxidative stress is associated with damage to intestinal epithelium, disturbed barrier function, and overactivation of conserved inflammatory pathways (e.g., NF-κB and JNK) [12–16]. Further, oxidative stress is a long-standing feature of aging and is linked to age-related diseases, including cancer [3, 17].

To respond to oxidative stress, metazoans employ numerous strategies to reduce host damage and maintain tissue and organismal homeostasis. ROS-sensitive signaling cascades, notably the NFE2L2 (Nrf2) and MAPK-signaling cascades, respond to oxidant burden by activating transcriptional programs which govern antioxidant capacity and stress adaptation (e.g., cell proliferation and cell death) [18–20]. Acting alongside these signaling cascades, the endolysosomal system degrades and recycles extracellular and intracellular material delivered through endocytosis, phagocytosis, and converging autophagic pathways [21, 22]. This system is activated in response to increased oxidative stress and inflammation and can serve as a homeostatic mediator to clear damaged proteins, inflammasome components, and invading pathogens [23–25]. Notably, IBD susceptibility is associated with polymorphisms in genes encoding Autophagy Related 16 Like 1 (ATG16L1) and Immunity-Related GTPase M (IRGM) [26–30] and lysosomal defects are linked to several of the hallmarks of aging [31, 32].

Lysosomes are the terminal organelles of the endolysosomal pathway and function to digest and recycle macromolecules [33]. Central to this system is luminal acidification. Cargo is delivered to endosomal compartments and maturation along the endocytic pathway involves progressive acidification of the lumen of endosomal compartments, from near neutral early endosomes to strongly acidic lysosomes [34, 35]. This acidification is generated by the highly conserved vacuolar ATPase (V-ATPase), a multi-subunit transmembrane proton pump that couples ATP hydrolysis to proton translocation across endomembranes [36, 37]. Because V-ATPase activity is continuously required, tunable, and energetically costly, endolysosomal pH is a well-regulated variable.

Increasingly, endosomal and lysosomal membranes have gained appreciation as regulatory hubs for signal transduction involved in cell death and survival, cell growth, autophagy, membrane damage and repair, microbe clearance, and cytokine signaling [38–42]. Through its control of luminal pH, the V-ATPase complex governs endosomal ligand-receptor dissociation, recycling, and lysosomal degradation [43, 44]. In particular, endolysosomal acidification and V-ATPase subunit activity have been linked directly to the activation of mTOR, Wnt, TGF-β, and Notch signaling pathways [43, 44]. Yet, whether endolysosomal acidification shapes the signaling pathways that govern cell fate and tissue homeostasis during intestinal oxidative stress has not been examined.

Here, we use *Drosophila* to define the role of endolysosomal acidification in intestinal injury and repair driven by oxidative stress. Given the established requirement for vesicular acidification in autophagic and lysosomal flux, we anticipated that impairing endolysosomal acidification would sensitize the intestine to oxidant-induced damage. Instead, we find that RNAi depletion of multiple V-ATPase complex subunits protects animals from lethality induced by either hydrogen peroxide or paraquat. These effects map to mature enterocytes (ECs), the absorptive cells of the *Drosophila* intestine. Oxidant exposure increases V-ATPase subunit expression and the abundance of acidified vesicles in the midgut, whereas EC-specific depletion of *Vha44* suppresses both this acidification and the cell death, barrier dysfunction, JNK activation, and IMD/NF-κB activation associated with oxidant induced injury. We find that overexpression of the MAP3K *Tak1*, a shared upstream node of the JNK and IMD/NF-κB cascades, sensitizes animals to oxidative stress and drives both pathways in a *Vha44*-dependent manner. Together, these findings identify endolysosomal acidification as an amplifier of the stress and innate immune signaling that the intestinal epithelium mounts in response to excessive oxidant burden.

## Materials and Methods

### *Drosophila* Strains and Culture

*Drosophila melanogaster* strains were maintained on standard diets at 25 °C with 12-h light/dark cycles (7 am–7 pm). Flies were maintained on standard cornmeal, agar, molasses medium at 25°C. The following lines were obtained from the Bloomington *Drosophila* Stock Center: *UAS-Vha44-IR* (BDSC #33884; validated in [45–47]), *UAS-Vha68-2-IR* (BDSC #34582; [45]), *UAS-VhaSFD-IR* (BDSC #40896; [46]), *UAS-cathD-IR* (BDSC #28978; [48]), *Vha55-lacZ* (BDSC #12128), *UAS-LAMP1-GFP* (BDSC #42714), *hid-GFP* (BDSC #50750), *puc-lacZ* (BDSC #98329), *UAS-Bsk* (BDSC #83654), *UAS-Bsk-IR* (BDSC # 57035), *Drs-GFP* (BDSC #55707), *Rel-GFP* (BDSC #43956), *UAS-Vha44* (BDSC #20140; [46]), *UAS-VhaSFD* (BDSC #15758; [46]). From Vienna *Drosophila* Stock Center: *UAS-Relish-KD* (VDRC #108469; [49]). Gifts from Rheinallt Jones w^1118^, *ß-tubulin-Gal4*, *escargot-Gal4*, *Myosin 1A-Gal4*, and *UAS-dTak1*. For Gal4/UAS experiments *ß-tub>+*, *esg>+*, and *Myo1A>+* indicates the respective Gal4 driver crossed to w^1118^ control flies.

### Drug Treatment and Lifespan Assays

Males of the specified genotypes were collected 2-5 days post-eclosion and placed on vehicle or drug treated food. Vehicle food was 5% sucrose in 1% agar. For injury models, paraquat (Sigma-Aldrich #856177) was dissolved at 10mM; hydrogen peroxide (VWR BDH7690-1) was mixed at 1% w/v; and DSS was dissolved at 2.5% w/v. For lifespan, the number of dead flies were counted daily. All flies used in an individual experiment were started on experimental food on the same day. Lifespan data was analyzed using the Kaplan-Meier method using GraphPad Software Prism version 11 and statistical comparisons were made using the log-rank (Mantel-Cox) test.

### Smurf Assay

Flies were fed 10mM PQ as in lifespan experiments and supplemented with 2.5% w/v Food Blue #1 (Sigma-Aldrich 3844-45-9). Each day flies were examined on a flypad. Once a fly became Smurf or died it was counted as Smurf or non-Smurf, as in [50]. Only those flies with extensive blue outside of their digestive tract were counted as Smurf. The Fisher’s exact test was then performed in GraphPad Software Prism version 11 to obtain P-values for the cumulative Smurf counts.

### Histology

Adult fly intestines were dissected at the times indicated (48hrs for H_2_O_2_ injury, 24hrs for PQ injury, and 24hrs for DSS injury unless stated otherwise), stained with standard immunohistochemical procedures, and mounted in Vectashield antifade mounting medium with DAPI (Vector Laboratories, H-1200). Antibodies used were mouse anti-β-Galactosidase (1:250, Promega Z378B), rabbit anti-cleaved Dcp-1 (Asp15) (1:100, Cell Signaling Technology, 9578S), rabbit anti-Phospho-histone H3 (1:500, Cell Signaling Technology, 9701S), and rabbit anti-activated p-JNK (1:100, Cell Signaling Technology 9251S). Fluorescently labeled secondary antibodies were from Jackson ImmunoResearch and used at 1:50. Intestines were analyzed by Olympus FV1000 confocal microscope system. All images were taken in the R4-R5 region of the midgut. For Lysotracker staining, adult intestines were dissected into Schneider’s *Drosophila* media (Gibco 21720024). Then stained with LysoTracker Red DND-99 (ThermoFisher Scientific, L7528) at 1μM in Schneider’s media for 10 minutes at room temperature in the dark. Intestines were washed 2x in Schneider’s media then nuclei were stained with Hoechst 33342 (Thermo Fisher Scientific) in Schneider’s for 5 minutes in the dark at room temperature, then mounted in *n*-propyl gallate (4% w/v in glycerol) and immediately analyzed.

### RNA Isolation and Quantification

For RNA extraction from intestines, we performed 3 replicates with 30 guts from each of the indicated genotypes. For whole animal RNA extraction, 10 adult males were used for each replicate. Fly intestines were homogenized in 100μl of TRIzol reagent (Invitrogen, Life Technologies) and adult whole flies in 500μL TRIzol reagent. We performed RNA extractions using the standard phenol-chloroform method and included a DNase treatment (Zymogen Research). cDNA conversion was performed using SuperScript-III RT kit (Thermo Fisher Scientific; 18080093) according to the manufacturer’s instructions, using 1 µg of extracted RNA. qPCR analysis was performed in triplicate using SYBR Green Master Mix (Qiagen, 204143) on a Biorad CFX Connect Real-Time System. qPCR analysis was performed using the ΔΔCT method and expression levels were normalized to *Rp49*. Statistical analysis was performed in GraphPad Prism 11 using a two-tailed Student’s t-test. Primers were ordered from Integrated DNA Technologies:

Rp49_Forward: AGCATACAGGCCCAAGATCG; Rp49_Reverse: TGTTGTCGATACCCTTGGGC;

Drs_Forward: TACTTGTTCGCCCTCTTCG; Drs_Reverse: GTATCTTCCGGACAGGCAGT;

DPTA_Forward: CCCGACGACATGACCATGAA; DPTA_Reverse: CCACTTTCCAGCTCGGTTCT;

ATTC_Forward: TCCGTATACCCAGCCACTGA; ATTC_Reverse: AGGCCGTGTCCATGATTGTT.

## Results

### V-ATPase subunits are required for oxidative-stress induced lethality

Vacuolar ATPases (V-ATPases) are highly conserved multi-subunit proton pumps that drive the progressive acidification of endosomes and lysosomes to shape signal transduction and enable cargo degradation [33, 38, 51]. To test the genetic requirement for *V-ATPase* in response to oxidative stress, we used the Gal4/UAS system to drive whole body RNAi mediated knockdown of three V-ATPase subunits and fed adult flies hydrogen peroxide (H_2_O_2_), a well-described oxidant [52]. Rather than sensitizing animals to oxidant, whole animal depletion of each of three V-ATPase subunits, including *Vha44* (ATP6V1C1 ortholog), *Vha68-2* (ATP6V1A ortholog), and *VhaSFD* (ATP6V1H ortholog), is protective as these *Vha*-depleted animals show increased survival compared to driver matched controls (ß-tub>+) (Fig 1A). Analysis of V-ATPase subunit expression using FlyAtlas 2 shows enhanced tissue expression in the intestine, Malpighian tubules, and salivary gland (Fig 1B, adapted from [53]). As the intestinal epithelium is directly exposed to oxidants in our feeding assay, we tested whether the protective effects of V-ATPase subunit depletion mapped to the intestinal epithelium. We used the intestine specific tissue drivers *escargot-Gal4* (*esg-Gal4*), which drives expression in intestinal stem cells (ISCs) and their enteroblast daughter cells, and *Myosin 1A-Gal4* (*Myo1A-Gal4*), which drives expression in mature absorptive enterocytes. While little effect was observed with *esg*-driven RNAi expression, we find that depletion of each of these V-ATPase subunits from mature ECs suppresses oxidative stress-induced lethality (Fig 1C-D). In parallel, we tested whether genetic depletion of *cathepsin D*, the principal lysosomal aspartic protease [48, 54], could impact organismal survival in the setting of oxidative stress. In contrast to V-ATPase subunit depletion, we find that depletion of *cathD* is not protective to H_2_O_2_ exposure (S1A-B Fig).

**Figure 1.**
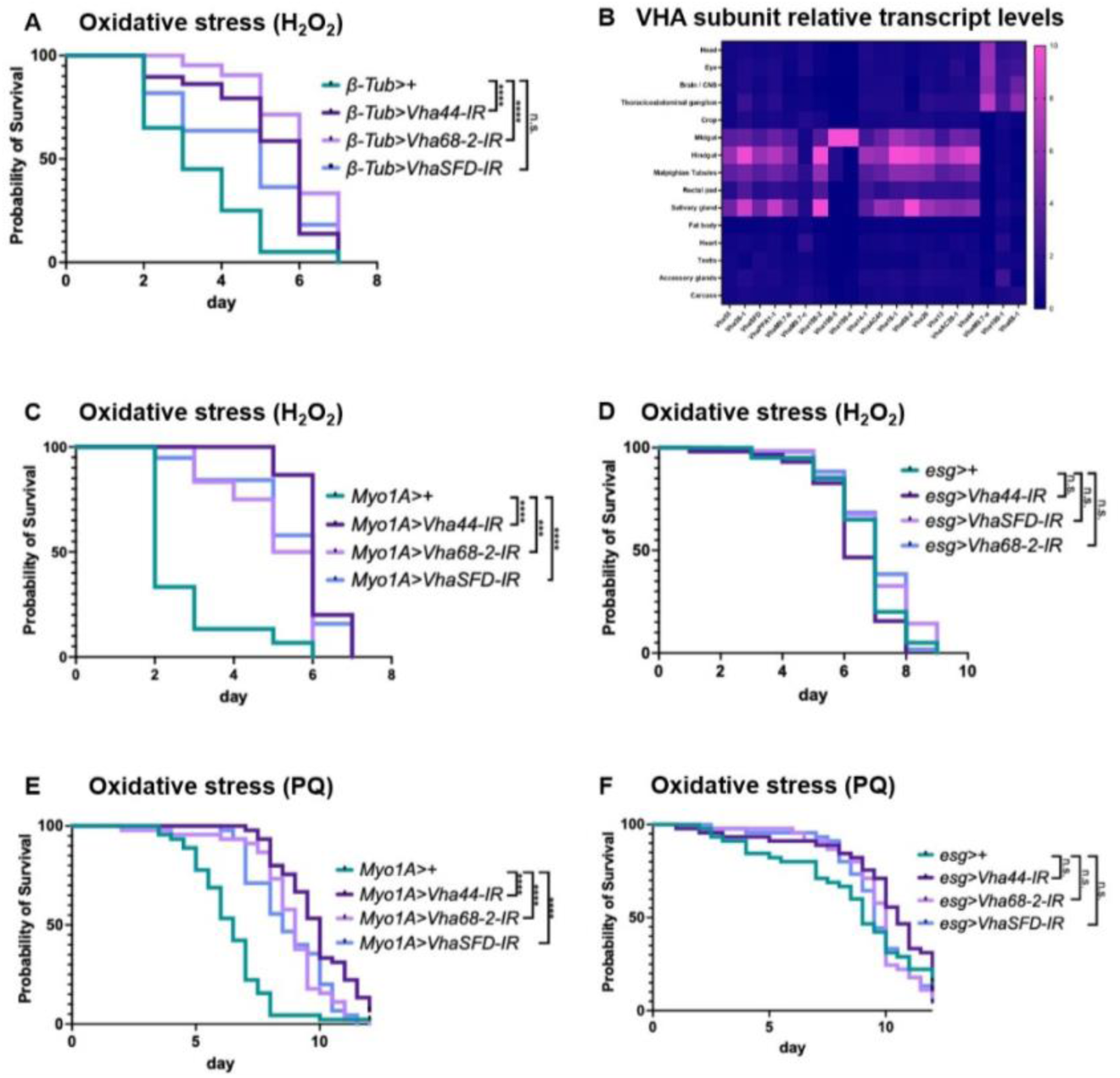
V-ATPase subunit depletion is protective against oxidative stress. Kaplan-Meier survival curves of adult male flies fed 1% H_2_O_2_ (A, C, D) or 10mM PQ (E, F), comparing RNAi depletion of the V-ATPase subunits *Vha44*, *Vha68-2*, or *VhaSFD* with driver matched controls (>+). A) Whole body driven depletion (*ß-tubulin-Gal4*) of flies fed 1% H_2_O_2_: *ß-tub>+* n=20, *ß-tub>Vha44-IR* n=28, *ß-tub>Vha68-2-IR* n=21, *ß-tub>VhaSFD-IR* n=11. B) Heat map of V-ATPase subunit transcript levels across adult tissues, generated from FlyAtlas 2 data [53]. C) Enterocyte-specific depletion (*Myo1A-Gal4*) of flies fed 1% H_2_O_2_: *Myo1A>+* n=45, *Myo1A>Vha44-IR* n=45, *Myo1A>Vha68-2-IR* n=42, *Myo1A>VhaSFD-IR* n=38. D) Progenitor-specific depletion (*esg-Gal4*; intestinal stem cells and enteroblasts) of flies fed 1% H_2_O_2_: *esg>+* n=40, *esg>Vha44-IR* n=58, *esg>Vha68-2-IR* n=60, *esg>VhaSFD-IR* n=49. E) Enterocyte-specific depletion of flies fed 10mM PQ: *Myo1A>+* n=44; *Myo1A>Vha44-IR* n=42; *Myo1A>Vha68-2-IR* n=45; *Myo1A>VhaSFD-IR* n=45. F) Progenitor-specific depletion of flies fed 10mM PQ: *esg>+* n=38; *esg>Vha44-IR* n=40; *esg>Vha68-2-IR* n=43; *esg>VhaSFD-IR* n=43. Statistics (A, C-F): log-rank (Mantel-Cox) test versus driver matched control; **** p<0.0001, *** p<0.001, n.s. p>0.05.

To test the generalizability of these findings, we examined a second, mechanistically distinct oxidant. Paraquat (PQ), a quaternary ammonium bipyridyl herbicide that induces oxidative stress in cells [55], produced the same result: EC-specific depletion of *Vha44*, *Vha68-2*, or *VhaSFD* protected animals from PQ-induced lethality (Fig 1E-F). Because increased ROS has been linked to aging across metazoans [3, 17, 56], and because intestinal barrier failure predicts mortality in *Drosophila* [57], we next asked whether V-ATPase depletion alters *Drosophila* lifespan in the absence of exogenous oxidative stress. Both whole-body and EC-specific depletion of *Vha44* extended lifespan relative to driver-matched controls on unchallenged food, while whole-body overexpression of V-ATPase subunits reduced lifespan (S1C-D Fig). These findings support V-ATPase as a general regulator of organismal survival to oxidative stress and implicate intestinal endosomal acidification as a regulator of lifespan and aging in metazoans.

### V-ATPase subunit expression, lysosomal reporter expression, and lysosomal acidification are upregulated in response to oxidative stress induced intestinal injury

V-ATPase subunit expression can be dynamically regulated in response to stress, including via the CLEAR (Coordinated Lysosomal Expression and Regulation) network [58–60]. We therefore asked whether V-ATPase expression is regulated by oxidative injury in the intestine, using the transcriptional reporter, *Vha55-* lacZ. Oxidant exposure significantly increased *Vha55-*lacZ expression in the R4-R5 region of the midgut (Fig 2A-C, E). This induction was not a generic injury response as we find *Vha55-*lacZ expression is unchanged following dextran sodium sulfate (DSS) administration, which damages the intestine through disruption of basement membrane organization [61] (Fig 2A-E).

**Figure 2.**
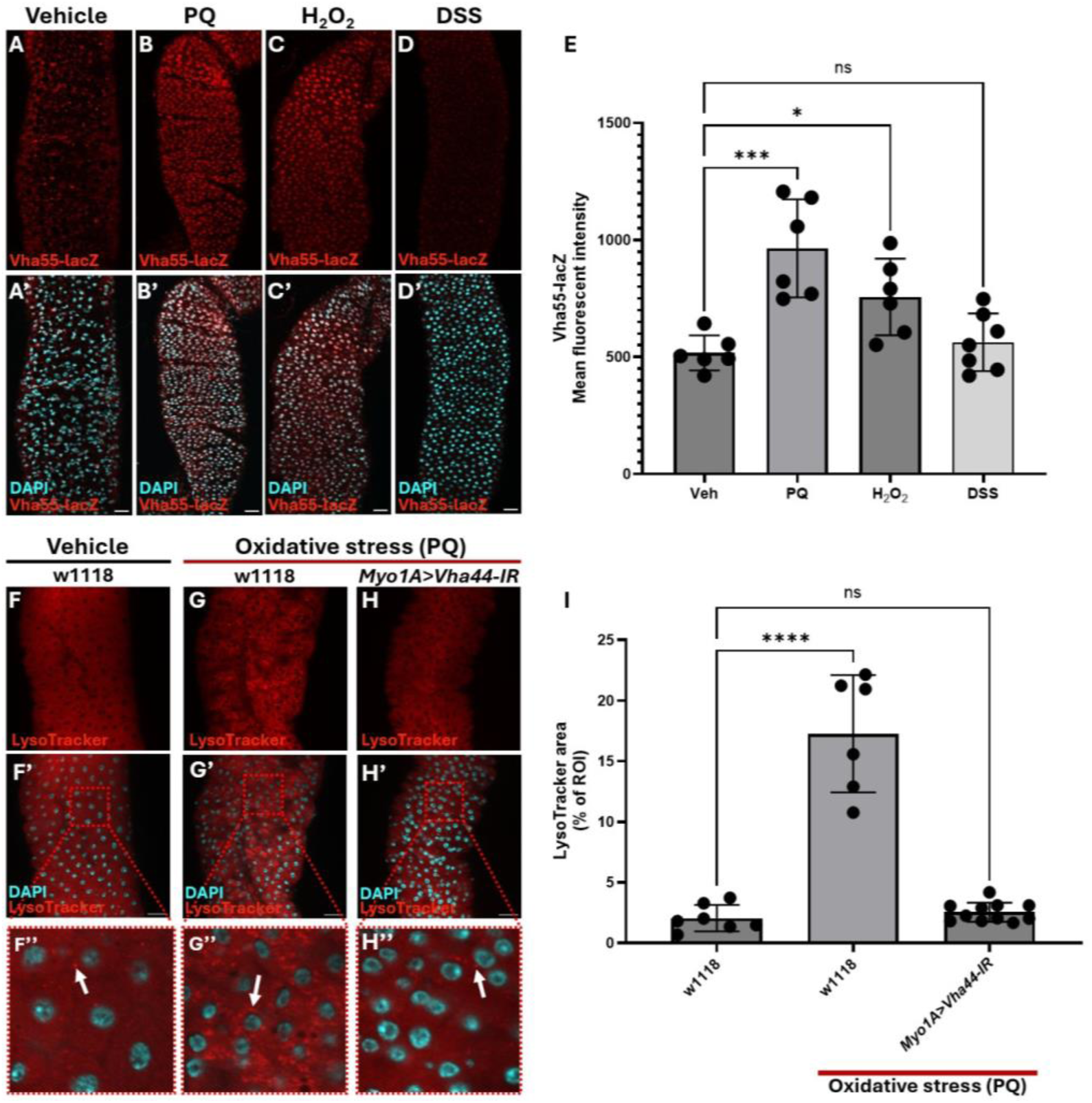
Oxidative injury induces V-ATPase subunit expression and endolysosomal acidification in the midgut. A-D’) Representative images of *Vha55*-lacZ transcriptional reporter in the R4-R5 region of the *Drosophila* midgut from adult males fed vehicle (A), 10mM PQ (B), 1% H_2_O_2_ (C), or 2.5% (w/v) DSS (D). Scale bars: 50 µm. E) Quantification of *Vha55*-lacZ mean fluorescent intensity. n=6-7 guts per condition. F-H’’) Representative images of LysoTracker Red staining in *w1118* animals fed vehicle (F) or PQ (G), and in *Myo1A>Vha44-IR* animals fed PQ (H). White arrows point to vesicular LysoTracker. Scale bars: 25 µm. I) Quantification of vesicular LysoTracker signal, as percentage of region of interest (ROI) area. Vehicle *w1118* n=7, PQ *w1118* n=6, PQ *Myo1A>Vha44-IR* n=11. Statistics (E, I): one-way ANOVA with Dunnett’s multiple comparisons test against the vehicle-fed control; **** p<0.0001, *** p<0.001, *p<0.05, n.s. p>0.05.

Consistent with a broader lysosomal program, expression of a Lysosomal-Associated Membrane Protein 1 reporter, LAMP1-GFP, was broadly increased in the midguts of oxidative-challenged animals (S2A-C Fig). Functionally, oxidant exposure also increased the abundance of acidified vesicular compartments, measured with the cell-permeable acidotropic dye LysoTracker Red, and this increase requires *Vha44* (Fig 2F-I). Together, these data indicate that oxidative injury elicits a coordinated increase in lysosomal biogenesis and endolysosomal acidification in the intestinal epithelium.

### V-ATPase subunits regulate intestinal injury and repair in *Drosophila*

Having established that oxidative injury induces V-ATPase expression and that V-ATPase depletion is protective, we next asked how *Vha44* depletion affects intestinal physiology. Maintenance of barrier function is a critical role of metazoan intestinal epithelium, and barrier failure is linked to systemic inflammation, morbidity, and mortality [57, 62, 63]. Using the Smurf assay [64], which measures intestinal permeability in *Drosophila*, we found that EC-specific *Vha44* depletion suppressed oxidant-induced barrier failure (Fig 3A-C).

**Figure 3.**
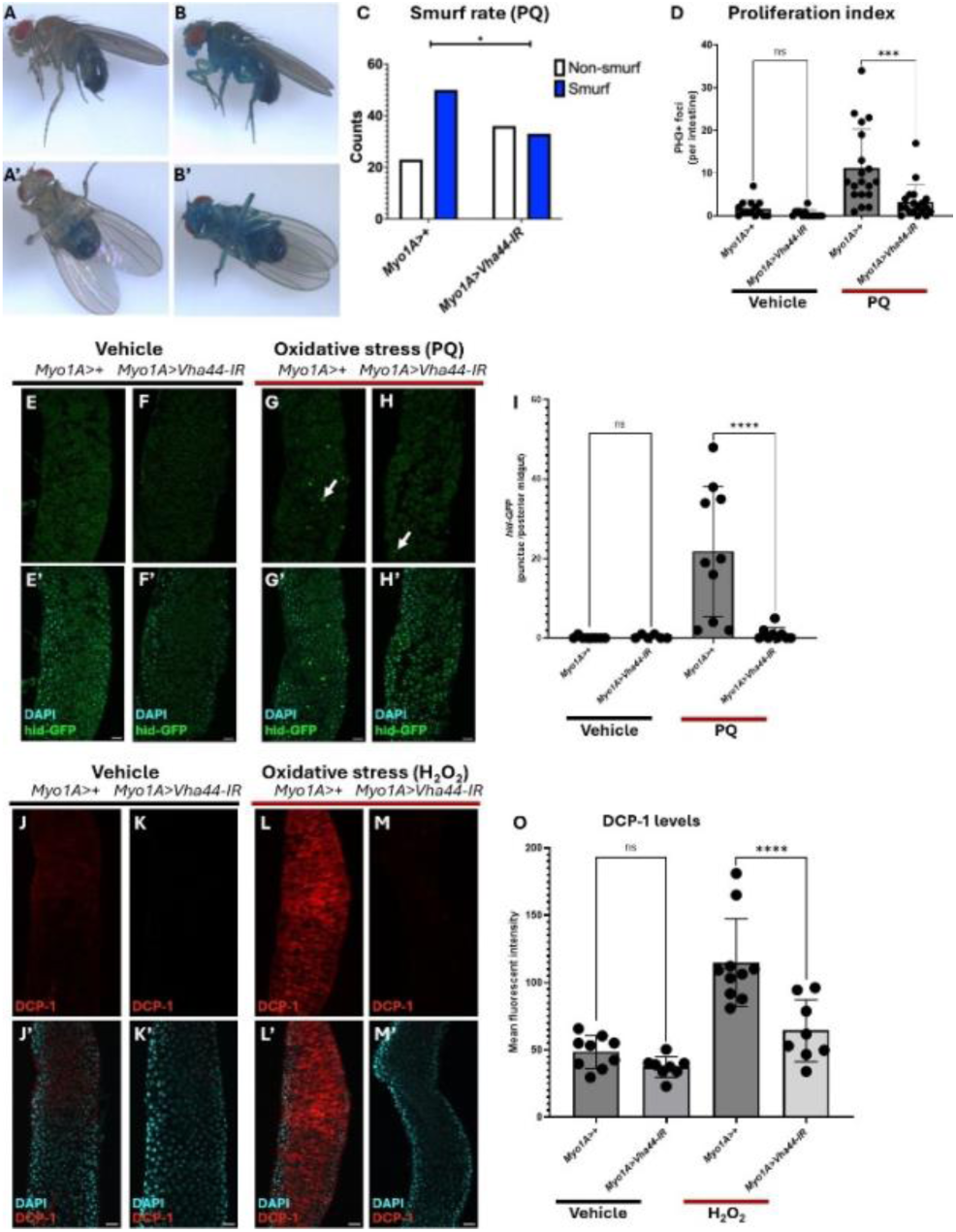
Enterocyte-specific *Vha44* depletion suppresses oxidant-induced intestinal barrier failure and cell death. A-B’) Representative images of *Myo1A>+* flies illustrating Smurf phenotype from side and ventral views of non-Smurf (A-A’) and Smurf (B-B’) flies, Smurf animals show dye distributed outside digestive tract. C) Number of *Myo1A>+* and *Myo1A>Vha44-IR* flies scored as Smurf or non-Smurf after 10mM PQ feeding; n=73 and n=69 respectively per genotype. Fisher’s exact test * p<0.05. D) Number of PH3+ foci per intestine. Vehicle: *Myo1A>+* n=14, *Myo1A>Vha44-IR* n=14; PQ: *Myo1A>+* n=19, *Myo1A>Vha44-IR* n=19. E-H’) Representative images of *hid*-GFP in R4-R5 midgut region of vehicle fed (E-F’) and PQ fed (G-H’) flies; white arrows indicate *hid*-GFP puncta. Scale bar: 25 µm. I) Number of *hid*-GFP puncta per high-power field. Vehicle: *Myo1A>+* n=8, *Myo1A>Vha44-IR* n=6; PQ: *Myo1A>+* n=9, *Myo1A>Vha44-IR* n=10. J-M’) Representative images of DCP-1 staining in R4-R5 midgut region of vehicle (J-K’) and 1% H_2_O_2_ (L-M’) fed flies. N) Mean fluorescence intensity of DCP-1 staining. Vehicle: *Myo1A>+* n=9, *Myo1A>Vha44-IR* n=8; H_2_O_2_: *Myo1A>+* n=10, *Myo1A>Vha44-IR* n=8. Statistics: D, I, N) One-way ANOVA with Šídák’s multiple comparisons test, **** p<0.0001, *** p<0.001, n.s. p>0.05.

Because midgut injury normally elicits compensatory proliferation [52, 65], we asked whether preserved barrier function reflected enhanced regeneration. We find that EC-specific *Vha44* depletion instead suppressed the proliferative response to oxidative stress (Fig 3D), consistent with reduced injury rather than increased repair. Since compensatory proliferation in fly epithelia is driven by apoptotic signals from dying cells [66, 67], this result predicted that cell death itself would be reduced. We therefore examined cell-death pathways. Oxidant exposure increased expression of the pro-apoptotic gene *hid*, reported by *hid*-GFP, and this induction was suppressed by EC-specific *Vha44* depletion (Fig 3E-I). In parallel, staining for the effector caspase Death caspase-1 (DCP-1) was increased following oxidative stress and likewise suppressed by *Vha44* depletion (Fig 3J-O, S3A-C). These findings indicate that *Vha44* depletion attenuates the enterocyte death and barrier failure that follow oxidant exposure, consistent with a model in which reduced endolysosomal acidification limits injury rather than accelerating repair.

### V-ATPase subunit expression is required for JNK up-regulation in response to oxidative stress

The c-Jun N-terminal Kinase (JNK) pathway is an evolutionarily conserved MAP kinase stress response pathway induced by oxidative stress in the intestine [65, 68, 69]. JNK output is not uniformly protective or destructive. Transient or moderate JNK activity can be cytoprotective and regenerative [66, 67, 70–73], while stronger and more sustained signaling drives apoptosis [74–78]. This threshold behavior is mediated in part by JNK-dependent transcription of the pro-apoptotic gene *hid* in *Drosophila* epithelia [75], making JNK a candidate mediator of the cell death we observed following oxidant exposure. Notably, ectopic *Vha44* expression in the wing disc is sufficient to activate JNK and induce apoptosis [79]. Consistent with this, oxidative stress increased pJNK staining in the midgut, and this increase was suppressed by EC-specific *Vha44* depletion (Fig 4A-E, S4A-C Fig). The JNK transcriptional reporter *puc-*lacZ was likewise strongly induced by oxidant exposure in a *Vha44*-dependent manner (Fig 4F-J, S4D-H Fig), as was *puc* transcript measured by qPCR on dissected intestines (Fig 4K).

**Figure 4.**
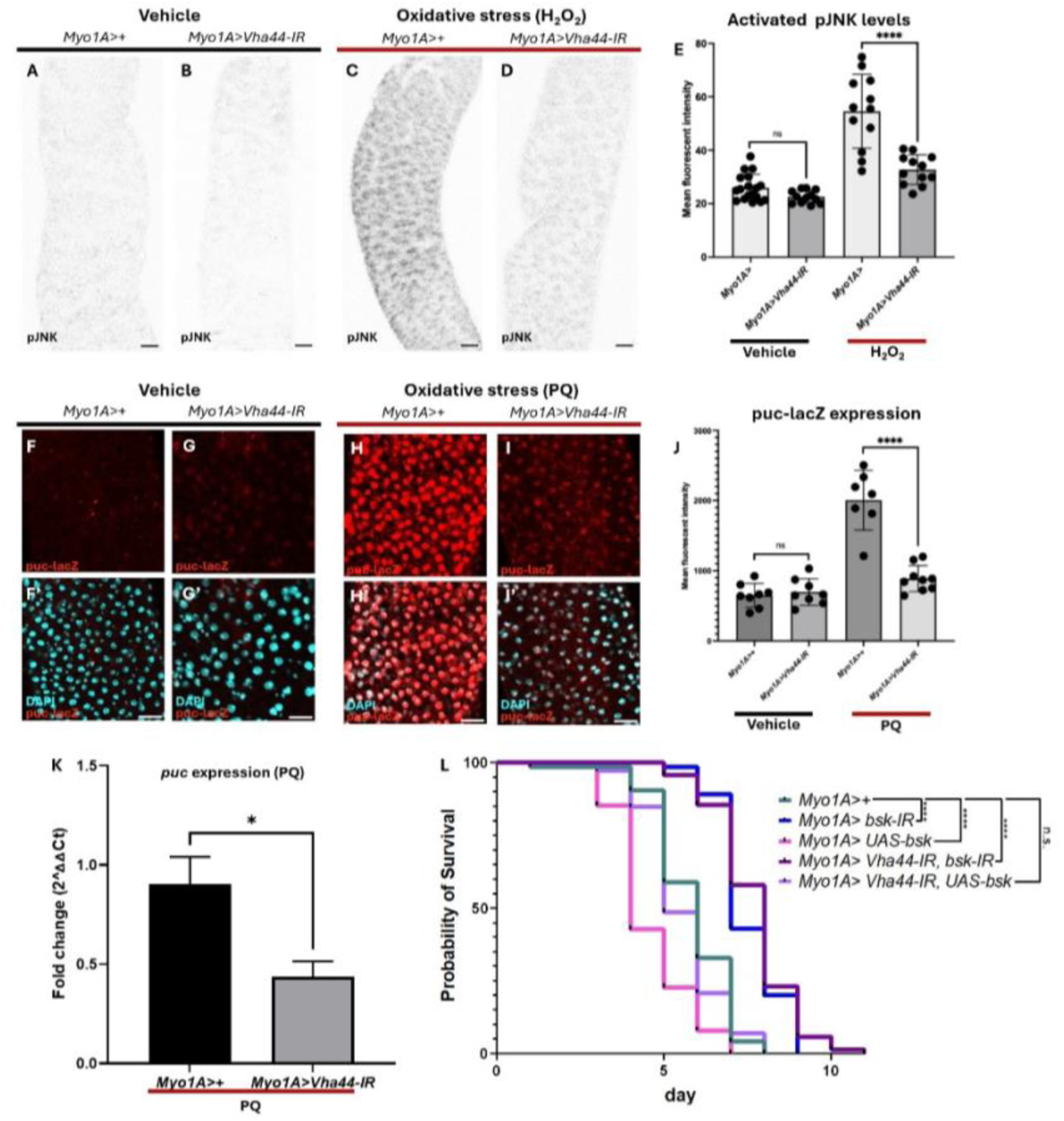
Oxidant-induced JNK activation in the midgut requires *Vha44*. A-D) Representative images of pJNK staining in R4-R5 midgut region of vehicle (A-B) and 1% H_2_O_2_ (C-D) fed flies. Scale bars: 25 µm. E) Mean fluorescence intensity of pJNK staining. Vehicle: *Myo1A>+* n=18, *Myo1A>Vha44-IR* n=12; H_2_O_2_: *Myo1A>+* n=12, *Myo1A>Vha44-IR* n=12. F-I’) Representative images of *puc*-lacZ stain in the R4-R5 midgut region of vehicle (F-G’) and PQ (H-I’) fed flies. Scale bars: 25 µm. J) Mean fluorescence intensity of *puc*-lacZ staining. Vehicle: *Myo1A>+* n=8, *Myo1A>Vha44-IR* n=8; PQ: *Myo1A>+* n=7, *Myo1A>Vha44-IR* n=9. K) Fold change in *puc* transcript in dissected midguts following PQ feeding, normalized to *Rp49*. n=3 per genotype. Student’s t-test, * p<0.05. L) Kaplan-Meier survival curve of flies fed PQ. *Myo1A>+* n=73; *Myo1A>bsk-IR* n=65; *Myo1A>UAS-bsk* n=75; *Myo1A>Vha44-IR, bsk-IR* n=69; *Myo1A>Vha44-IR, UAS-bsk* n=72. Log-rank (Mantel-Cox) test, **** p<0.0001, n.s. p>0.05. Statistics (E,J): one-way ANOVA with Šídák’s multiple comparisons test; **** p<0.0001, n.s. p>0.05.

To test whether JNK activity is functionally coupled to oxidant-induced lethality, we manipulated *basket* (*bsk*), the *Drosophila* JNK ortholog, in ECs. EC-specific *bsk* depletion suppressed oxidant-induced lethality, whereas *bsk* overexpression enhanced it. Co-depletion of *Vha44* reversed the sensitization conferred by *bsk* overexpression (Fig 4L). Because overexpressed Bsk still requires phosphorylation by upstream kinases to become active, this result is consistent with *Vha44* acting at or above the level of JNK activation rather than downstream of it. Together with the suppression of hid-GFP and DCP-1 (Fig 3E-O), these data place V-ATPase-dependent acidification upstream of JNK driven pro-apoptotic signaling following oxidant injury in ECs.

### V-ATPase subunit expression is required for IMD upregulation in response to oxidative stress

JNK signaling engages in extensive crosstalk with innate immune pathways across metazoans, most directly through the shared MAP3K Tak1, which branches to both JNK and NF-κB pathways in *Drosophila* [80–82]. We therefore asked whether the IMD/NF-κB arm is likewise engaged during oxidative injury and also requires V-ATPase activity. EC-specific depletion of *Relish*, the IMD pathway NF-κB ortholog, significantly increased survival following PQ exposure (Fig 5A), indicating that oxidant-induced lethality depends on IMD pathway output in ECs. Consistent with pathway engagement, a *Relish* reporter, *Rel*-GFP, was induced in the midgut following oxidant exposure using H_2_O_2_ or, to a lesser extent, paraquat (Fig 5B-E, S5A-C Fig) and H_2_O_2_ induced expression was suppressed by *Vha44* depletion (Fig 5B-E).

**Figure 5.**
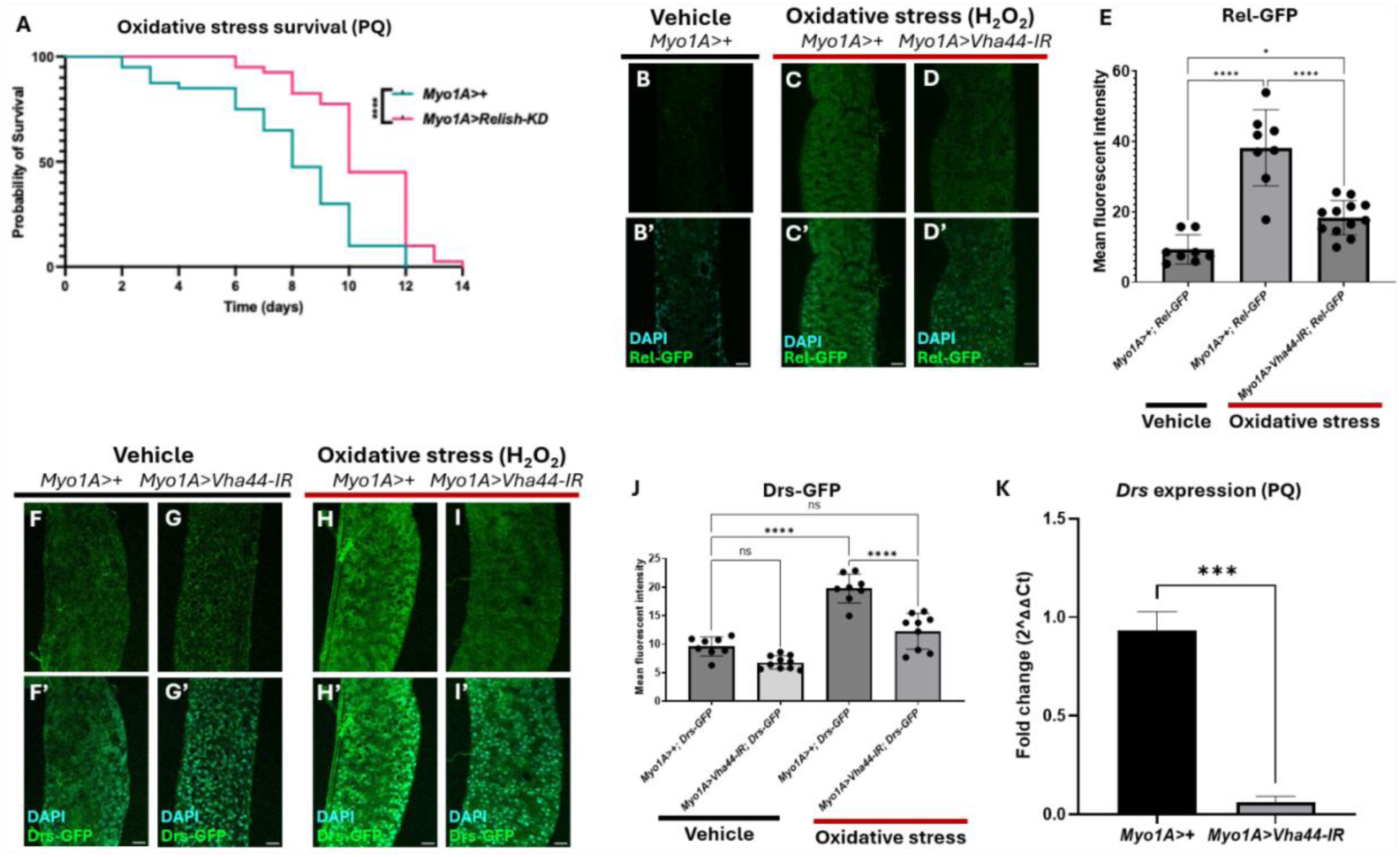
Oxidant-induced IMD/NF-κB-activation in the intestine requires *Vha44*. A) Kaplan-Meier survival curve of flies fed 10mM PQ. *Myo1A>+* n=45; *Myo1A>Relish-KD* n=45. Logrank (Mantel-Cox) test, *** p<0.001. B-D’) Representative images of *Relish*-GFP reporter in the R4-R5 midgut region of vehicle (B-B’) and 1% H_2_O_2_ (C-D’) fed flies. Scale bars: 25 µm. E) Mean fluorescence intensity of *Relish*-GFP. Vehicle: *Myo1A>+; Rel-GFP* n=8; H_2_O_2_: *Myo1A>+; Rel-GFP* n=8; *Myo1A>Vha44-IR; Rel-GFP* n=12. F-I’) Representative images of the *Drs*-GFP reporter in the R4-R5 midgut region of vehicle (F-G’) and 1% H_2_O_2_ (H-I’) fed flies. Scale bars: 25 µm. J) Mean fluorescence intensity of *Drs*-GFP. Vehicle: *Myo1A>+; Drs-GFP* n=8, *Myo1A>Vha44-IR; Drs-GFP* n=10; H_2_O_2_: *Myo1A>+; Drs-GFP* n=8, *Myo1A>Vha44-IR; Drs-GFP* n=9. K) Fold-change in *Drs* transcript in dissected midguts following PQ feeding, normalized to *Rp49*. n=3 per genotype. Student’s t-test, *** p<0.001. Statistics (E, J): one-way ANOVA with Tukey’s test for multiple comparisons. **** p<0.0001, * p<0.05.

To assess transcriptional output of the pathway, we used an *in vivo* reporter for the antimicrobial peptide (AMP) *Drosomycin*, *Drs*-GFP [83]. Although *Drosomycin* is a canonical Toll target during the systemic response, its local induction in barrier epithelia—including in the gut—is instead controlled by the IMD pathway [84, 85]. *Drs*-GFP was induced in a *Vha44*-dependent manner following either H_2_O_2_ (Fig 5F-J) or PQ (S5D-F Fig). In parallel, qPCR on dissected intestines confirmed *Vha44*-dependent induction of *Drs* transcript (Fig 5K).

### V-ATPase subunit expression is required for Tak1-induced oxidative stress response

The MAP3 kinase Tak1 acts upstream of both JNK and IMD signaling [86–88]. Furthermore, in mammalian cells, TAK1 is recruited to damaged endolysosomal membranes, where it activates downstream NF-κB and AMPK responses [89]. Tak1 therefore represents a plausible point at which endolysosomal state could be converted into stress and inflammatory pathway output, and we asked whether *Tak1* overexpression modifies the intestinal response to oxidative stress. EC-specific overexpression of *Tak1* sensitized flies to oxidative stress, and co-depletion of *Vha44* rescued this sensitization (Fig 6A). *Tak1* overexpression alone did not increase pJNK staining in unchallenged animals, but produced strong pJNK induction following oxidant exposure, consistent with Tak1 acting as a signal integrator rather than a constitutive activator. This oxidant-dependent pJNK induction was suppressed by *Vha44* depletion (Fig 6B-F).

**Figure 6.**
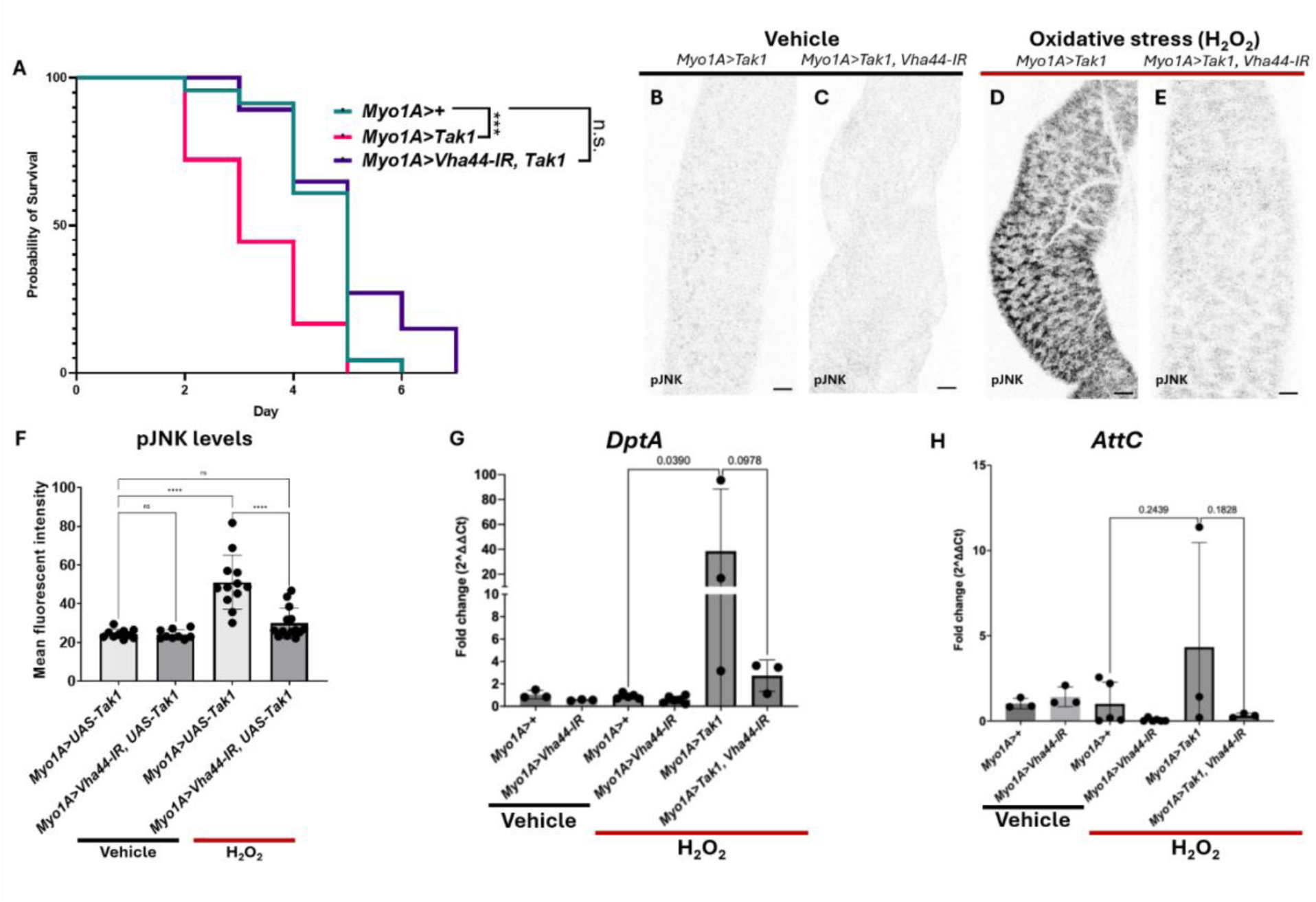
*Vha44* is required for Tak1-driven sensitization to oxidative stress and systemic IMD induction. A) Kaplan-Meier survival curve of flies fed 1% H_2_O_2_. *Myo1A>+* n=23; *Myo1A>UAS-Tak1* n=18; *Myo1A>Vha44-IR; UAS-Tak1* n=36. Log-rank (Mantel-Cox) test, *** p<0.001, n.s. p>0.05. B-E) Representative images of pJNK staining in R4-R5 midgut region of vehicle (B, C) and 1% H_2_O_2_ (D, E) fed flies. Scale bars: 25 µm. F) Mean fluorescence intensity of pJNK staining. Vehicle: *Myo1A>UAS-Tak1* n=10, *Myo1A>Vha44-IR; UAS-Tak1* n=9; H_2_O_2_: *Myo1A>UAS-Tak1* n=12, *Myo1A>Vha44-IR; UAS-Tak1* n=12. One-way ANOVA with Šídák’s multiple comparisons test, **** p<0.0001, n.s. p>0.05. G-H) Fold change in *DptA* (G) and *AttC* (H) transcripts from whole body of adult males fed vehicle or 1% H_2_O_2_, normalized to *Rp49*. n=3-5 for each genotype. One-way ANOVA with Šídák’s multiple comparisons test.

Finally, we asked whether the local intestinal response propagates to a systemic humoral response. The fat body is the dominant source of circulating AMP transcript [90, 91], so whole-animal measurements primarily report systemic rather than intestinal induction. At 48 hours post-H_2_O_2_ feeding, neither *DptA* nor *AttC* was elevated relative to vehicle-fed controls (Fig 6G-H), indicating that oxidant-induced IMD activation at this timepoint remains confined to the intestinal epithelium and does not elicit a detectable systemic response. However, in animals with EC-specific *Tak1* overexpression we see increased systemic *DptA* and a trend toward increased *AttC* and this increase was reduced by *Vha44* depletion (Fig 6G-H). Taken together, these results identify Tak1 as a point at which V-ATPase activity is required to convert oxidative stress into JNK and IMD/NF-κB output, consistent with the acidification dependence of both pathways observed above.

## Discussion

In the face of diverse endogenous and exogenous insults—including xenobiotics, metabolic disturbances, microbial challenges, and excessive inflammation—the intestinal epithelium must sense and calibrate its response to oxidative stress to preserve tissue homeostasis. Here we identify a previously unrecognized requirement for endolysosomal acidification in the intestinal inflammatory response to oxidative injury. Oxidant exposure rapidly increases V-ATPase subunit expression, lysosomal membrane reporter expression, and the quantity of acidified endolysosomal vesicles in the midgut. Yet preventing this acidification by depleting V-ATPase subunits protects animals across multiple oxidative injury models. At the cellular level, protection maps to mature enterocytes and is accompanied by reduced pro-apoptotic signaling and attenuated activation of the JNK and IMD/NF-κB pathways. Genetically, these effects require *Vha44* downstream of the MAP3K Tak1, a kinase recruited to damaged endolysosomal membranes in mammalian cells. Together our findings indicate that endolysosomal acidification regulates the magnitude of stress and innate immune signaling in the intestinal epithelium, linking the state of endolysosomal compartments to whether oxidative exposure is tolerated or resolved through cell death.

Recent work indicates that the restriction to mature enterocytes is meaningful. A genome-wide RNAi screen in adult midgut progenitors recovered V-ATPase subunits, including *Vha44* and *Vha68-2*, as required for ISC proliferation and differentiation under physiologic conditions, and for regenerative proliferation following ER stress. In progenitors under physiologic conditions, V-ATPase loss drives autophagy induction and ectopic Notch activation, leading to precocious differentiation at the expense of stem cell fate [92]. Yet even as stem cell function is compromised, progenitor V-ATPase depletion has no appreciable effect on barrier integrity, consistent with the progenitor-independent mechanism our data support [92]. Together, these observations indicate that V-ATPase serves functionally distinct roles in the two compartments of the midgut: maintenance of stem cell fate in progenitors, and amplification of stress and inflammatory signaling in mature ECs. Consistent with a broader role in age-associated intestinal decline, progenitor V-ATPase depletion has also been reported to block the loss of intestinal homeostasis otherwise seen in aged animals [92], paralleling the lifespan extension we observe with both whole body and EC-specific depletion (S1C-D Fig).

This work adds to a growing body of evidence implicating endolysosomal acidification, and the V-ATPase that generates it, as effectors of signal transduction in metazoans (reviewed in [43, 93, 94]). V-ATPase activity is required for amino acid–dependent activation of mTORC1 at the lysosomal surface, acting through amino acid–sensitive interactions with the Ragulator–Rag GTPase complex [95, 96]. In *Drosophila*, V-ATPase-dependent acidification is required not only for lysosomal turnover of the Notch receptor but for its activating cleavage, and for Wnt/Wingless signal reception—establishing that acidification can license signaling output rather than merely terminate it [97–99]. Our findings extend this principle to stress and innate immune signaling, where oxidant-driven JNK and IMD/NF-κB activation in the intestinal epithelium requires V-ATPase generated endolysosomal acidification.

These results also build upon prior work linking V-ATPase activity to cell death and cell competition in *Drosophila* epithelia [45, 79]. Ectopic *Vha44* expression in the wing disc activates JNK and induces apoptosis and tissue invasion [79], the gain-of-function converse of our loss-of-function result in the intestine. In the eye disc, V-ATPase subunits are autonomously required for elimination of “loser” cells through NF-κB-dependent induction of the pro-apoptotic gene *hid* and cooperative JNK-pathway signaling that promote loser cell death [45]. Our studies extend this cell death axis to the intestinal epithelium in the setting of oxidative injury. In this setting, we believe endolysosomal acidification serves as a regulator of cytoprotective and apoptotic programs needed to promote tissue homeostasis and organismal survival downstream of oxidative damage.

A central question raised by our results is why the intestine mounts a response, increased endolysosomal acidification, that proves deleterious. Release of ROS is a primary antimicrobial response in the intestinal epithelium, killing pathogens at the cost of host tissue damage [5, 8, 9, 100]. We suggest that endolysosomal amplification of cell stress and innate immune signaling is normally pro-adaptive, tuning the epithelial response to the magnitude of the microbial challenge, but that under excess oxidant burden this amplification contributes to additional damage to host tissue. An analogous principle operates in ECs, where autophagy lowers epithelial sensitivity to DUOX-derived ROS such that autophagy-deficient cells respond to substantially lower oxidant concentrations [101]. Endomembrane state therefore determines how strongly the epithelium responds to oxidant burden.

The control of innate immune signaling we describe appears to be local: whole-animal *DptA* and *AttC* transcripts were unchanged at 48 h in both control and Vha44-depleted animals, consistent with systemic AMP induction in *Drosophila* being contingent on intestinal barrier failure, which is not yet widespread at this timepoint [57, 71]. That EC-specific *Tak1* overexpression produced a trend toward elevated systemic AMP expression alongside accelerated oxidant-induced lethality, both reversed by *Vha44* co-depletion, is most consistent with systemic induction arising as a downstream consequence of barrier compromise rather than as a primary response, though we did not assess barrier integrity directly in these animals.

Acidification may act on complexes shared by both JNK and IMD/NF-κB pathways. Tak1 is an attractive candidate component: it is the shared MAP3K at which JNK and IMD signaling diverge, it is recruited to damaged endolysosomal membranes through TAB2/TAB3 recognition of K63-linked polyubiquitin in mammalian cells [89, 102, 103], and endosomal sorting defects activate Tak1-dependent JNK signaling in *Drosophila* [104]. Consistent with this we find that *Tak1*-driven JNK and IMD activation requires *Vha44*. Notably, *Tak1* overexpression alone did not induce pJNK in unchallenged animals, indicating that Tak1 requires a coincident oxidant-derived input, indicative of Tak1 as a signal integrator rather than a constitutive activator. Our conclusions regarding Tak1 rest on genetic epistasis; determining whether Tak1 localizes to endolysosomal membranes in ECs, and whether that localization is acidification-dependent, is a prime question this work opens.

We note that impaired acidification and impaired degradative flux are not mutually exclusive and our assessment of acidification relies on Lysotracker, which reports dye accumulation rather than luminal pH directly. However, genetic depletion of V-ATPase subunits in *Drosophila*, including subunits spanning both the V1 and V0 sectors, impairs vesicular acidification—producing LysoTracker negative compartments and preventing quenching of tandem-tagged Atg8a—while leaving membrane fusion intact [105]. The same dissociation is observed in mammalian cells, where inhibition or knockdown of V0 sector subunits reduces acidity without disrupting autophagosome-lysosome fusion [106, 107]. In the intestinal epithelium, V-ATPase-depleted progenitors likewise fail to quench tandem-tagged Atg8a, though the authors interpret this as reflecting impaired autophagic flux rather than reduced acidification [92]. That depletion of *cathD* does not phenocopy V-ATPase loss, together with the observation that Cathepsin depletion likewise fails to phenocopy V-ATPase depletion in other *Drosophila* tissues [105], further argues that loss of proteolytic capacity alone is insufficient to account for the phenotypes described. We therefore interpret the reduction in acidified vesicular compartments we observe (Fig 2F-I) as reflecting impaired acidification, though ratiometric measurement of luminal pH in ECs would be required to establish the magnitude of the shift.

These findings may bear on chronic inflammatory disease of the intestine, where epithelial damage in Crohn’s disease and ulcerative colitis arises substantially from the host response and IBD susceptibility loci converge on endolysosomal and autophagic genes including *ATG16L1* and *IRGM* [26–30]. Whether acidification represents a viable therapeutic target nonetheless requires care, because the direction of the V-ATPase requirement is cell-type and context dependent: even within the *Drosophila* intestine, depletion in ECs is protective whereas depletion in progenitors compromises stem cell fate [92]. Furthermore, the canonical tools are imperfect. Bafilomycin A1 has not been developed clinically owing to systemic toxicity [108, 109], and although chloroquine and hydroxychloroquine raise endolysosomal pH and dampen endosomal Toll-like receptor signaling, their dose-response relationship remains poorly defined [110, 111]. Nonetheless these tools provide proof of principle that luminal pH can be modulated, and this modulation can reduce inflammatory output rather than simply arrest degradation. This acidification-dependent node may warrant attention as a point of intervention under conditions of excess epithelial oxidant burden.

In conclusion, we show that V-ATPase activity and endolysosomal acidification are required for the intestinal epithelium to mount stress and innate immune signaling in response to oxidative injury, and that removing this requirement is protective. These findings identify endolysosomal pH as a tunable input to inflammatory signaling.

## Author Contributions

All authors listed made a substantial, direct and intellectual contribution to the work, and approved it for publication.

## Footnotes

This work was supported by the Emory University Integrated Cellular Imaging Core Facility (RRID:SCR_023534) Stocks obtained from the Bloomington Drosophila Stock Center (NIH P40OD018537) and the Vienna Drosophila Resource Center (VDRC, www.vdrc.at) were used in this study.

## Data Availability

All data are within the manuscript and its supporting information files

