## SupplementalFigures for "Endolysosomal acidification amplifies JNK and IMD/NF-κB signaling and exacerbates oxidative injury in the *Drosophila* intestine"

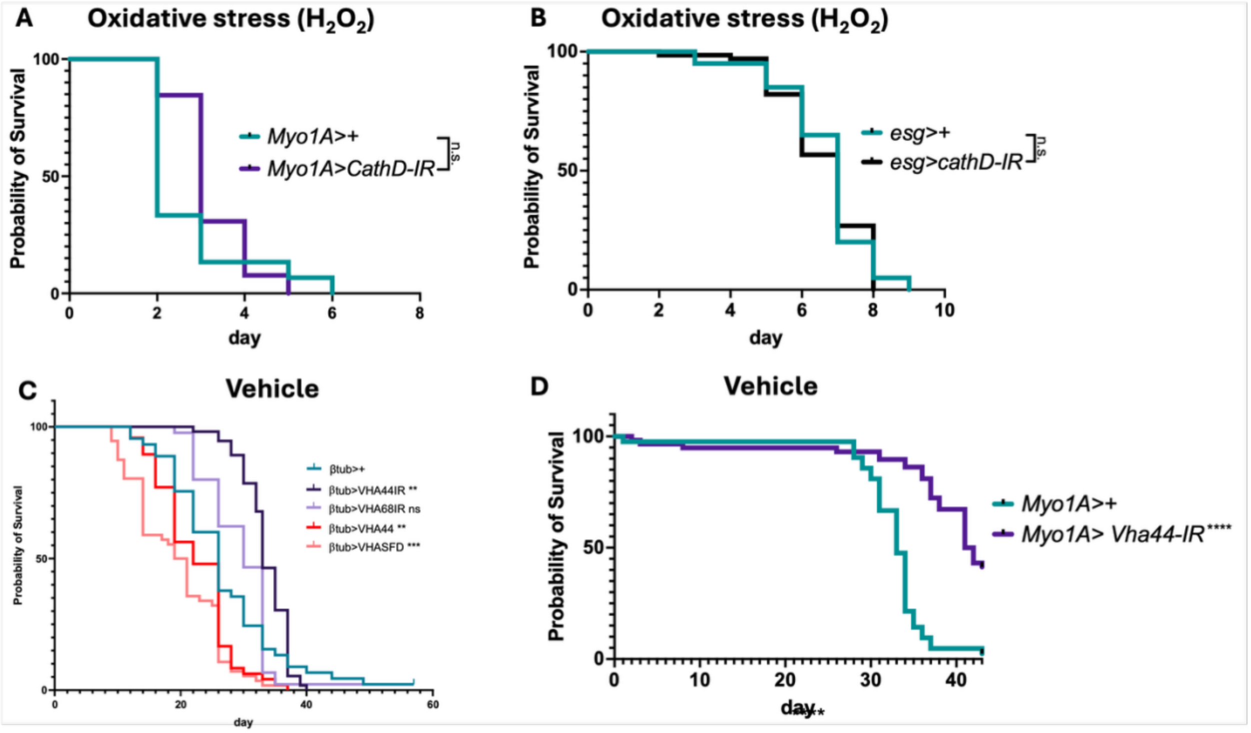
A
Oxidative stress ($H_2O_2$)
Probability of Survival
100
50
0
Myo1A>+
Myo1A>CathD-IR
n.s.
0
2
4
6
8
day
B
Oxidative stress ($H_2O_2$)
Probability of Survival
100
50
0
esg>+
esg>cathD-IR
n.s.
0
2
4
6
8
10
day
C
Vehicle
Probability of Survival
100
50
0
βtub>+
βtub>VHA44IR **
βtub>VHA68IR ns
βtub>VHA44 **
βtub>VHASFD ***
0
20
40
60
day
D
Vehicle
Probability of Survival
100
50
0
Myo1A>+
Myo1A> Vha44-IR****
0
10
20
30
40
day****

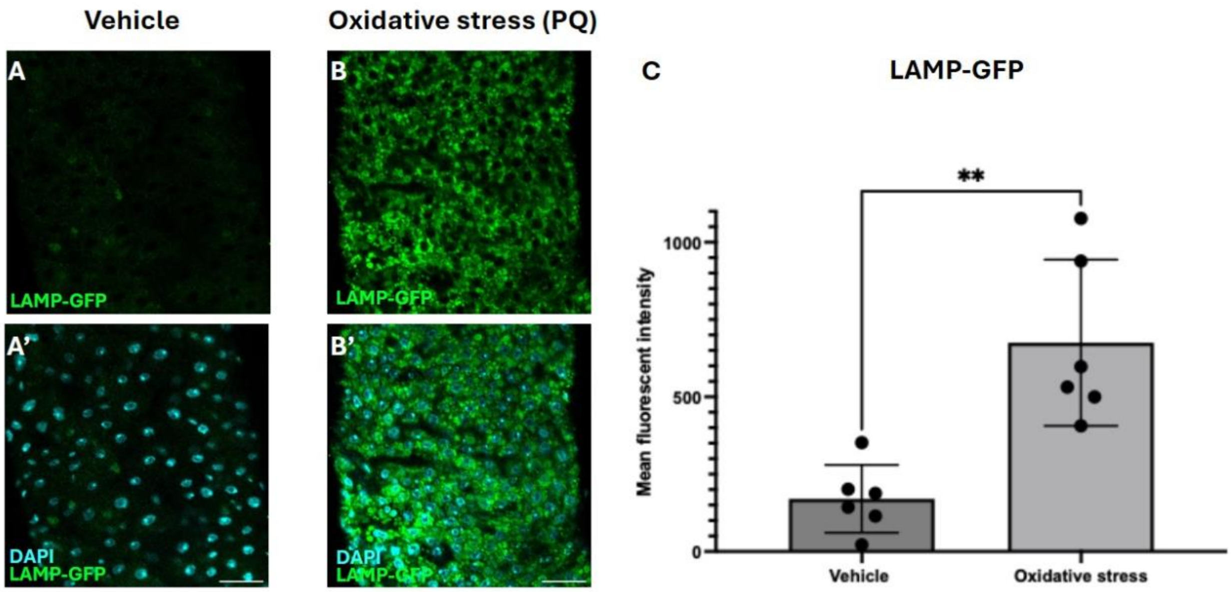
Vehicle
Oxidative stress (PQ)
A
LAMP-GFP
B
LAMP-GFP
A'
DAPI
LAMP-GFP
B'
DAPI
LAMP-GFP
C
LAMP-GFP
**
Mean fluorescent intensity
1000
500
0
Vehicle
Oxidative stress

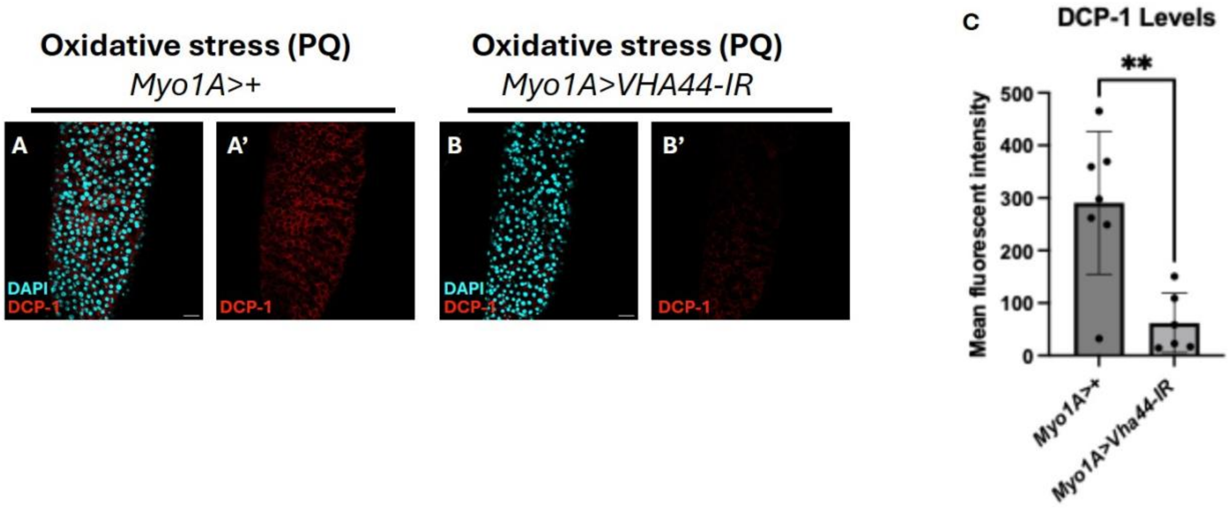
Oxidative stress (PQ)
Myo1A>+
Oxidative stress (PQ)
Myo1A>VHA44-IR
A
A'
B
B'
DAPI
DCP-1
DCP-1
DAPI
DCP-1
DCP-1
C
DCP-1 Levels
**
Mean fluorescent intensity
500
400
300
200
100
0
Myo1A>+
Myo1A>Vha44-IR

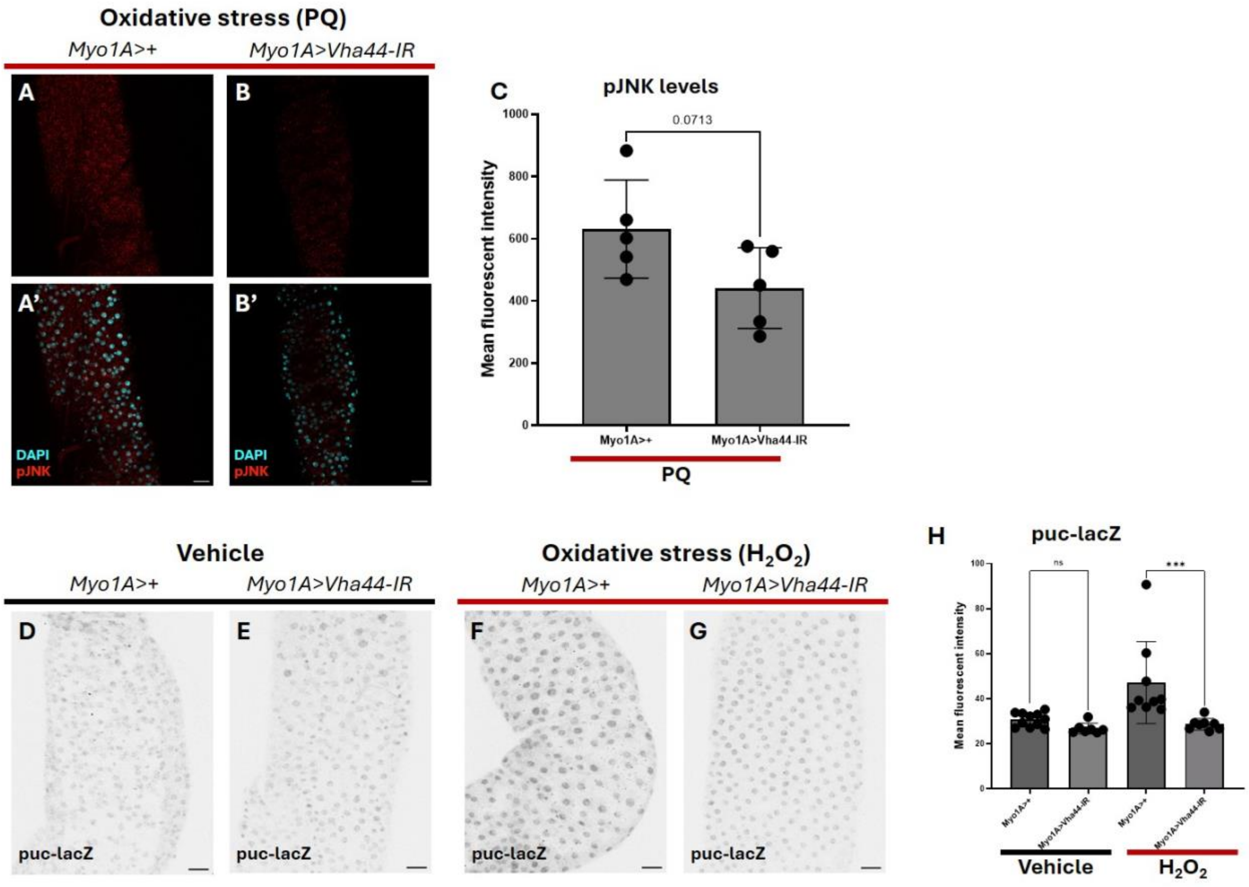

Oxidative stress (PQ)
Myo1A>+
Myo1A>Vha44-IR
A
B
A'
B'
DAPI
pJNK
DAPI
pJNK
C
pJNK levels
0.0713
Mean fluorescent intensity
1000
800
600
400
200
0
Myo1A>+
Myo1A>Vha44-IR
PQ
Vehicle
Myo1A>+
Myo1A>Vha44-IR
Oxidative stress (H2O2)
Myo1A>+
Myo1A>Vha44-IR
D
E
F
G
puc-lacZ
puc-lacZ
puc-lacZ
puc-lacZ
H
puc-lacZ
ns
***
Mean fluorescent intensity
100
80
60
40
20
0
Myo1A>+
Myo1A>Vha44-IR
Myo1A>+
Myo1A>Vha44-IR
Vehicle
H2O2

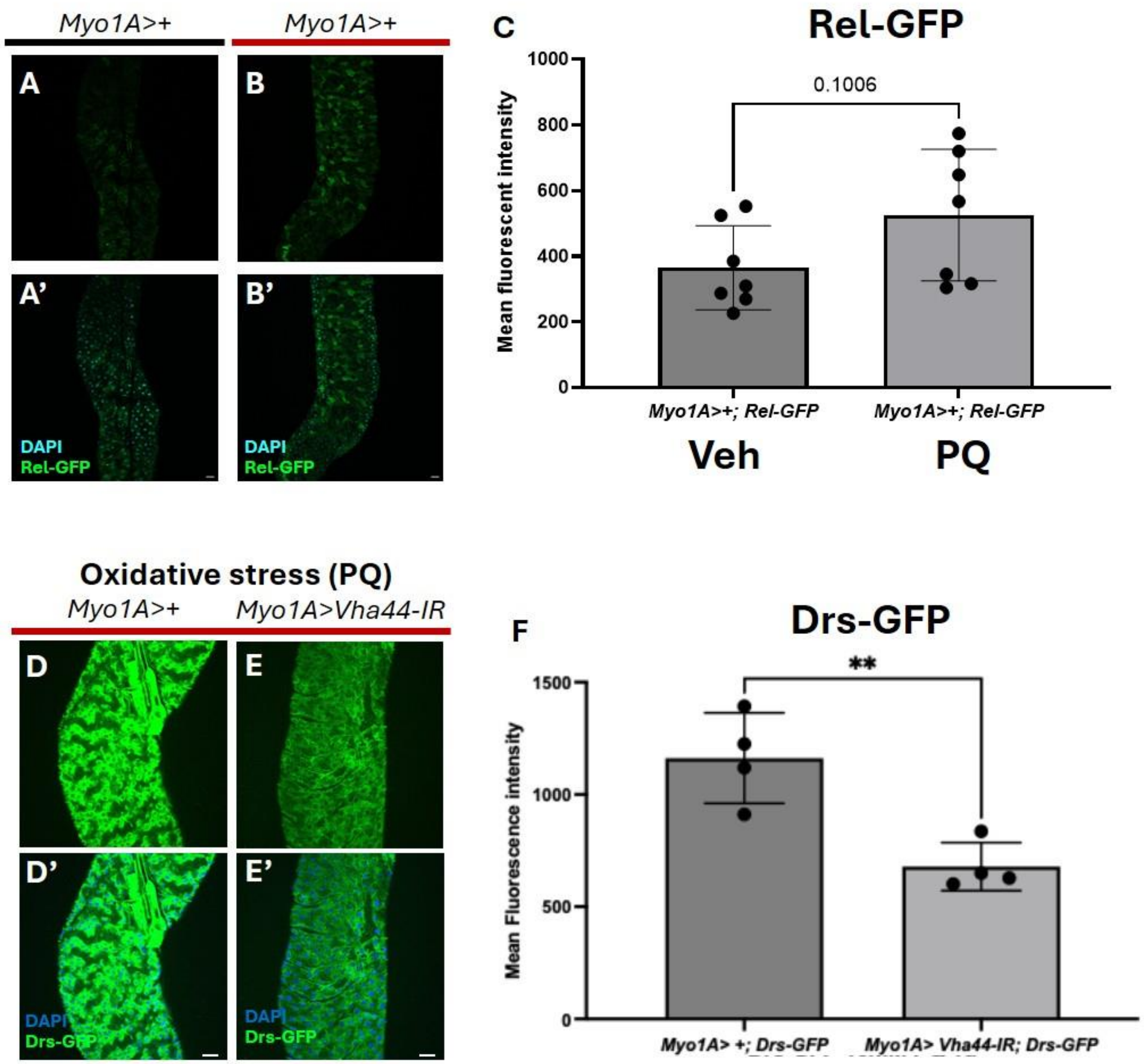

Myo1A>+
Myo1A>+
A
B
A’
B’
DAPI
Rel-GFP
DAPI
Rel-GFP
C
Rel-GFP
0.1006
Mean fluorescent intensity
1000
800
600
400
200
0
Myo1A>+; Rel-GFP
Myo1A>+; Rel-GFP
Veh
PQ
Oxidative stress (PQ)
Myo1A>+
Myo1A>Vha44-IR
D
E
D’
E’
DAPI
Drs-GFP
DAPI
Drs-GFP
F
Drs-GFP
**
Mean Fluorescence intensity
1500
1000
500
0
Myo1A> +; Drs-GFP
Myo1A> Vha44-IR; Drs-GFP

**S1 Fig. *Cathepsin D* depletion does not phenocopy V-ATPase loss, and V-ATPase depletion extends lifespan.** A, B) Kaplan–Meier survival curves of flies fed 1% $H_2O_2$ following cathD RNAi depletion driven in enterocytes (A; *Myo1A-Gal4*) or intestinal progenitors (B; *esg-Gal4*). A) *Myo1A*>+ n=45, *Myo1A >cathD-IR* n=39. B) *esg*>+ n=40, *esg >cathD-IR* n=67. C) Whole-body manipulation (*β-tubulin-Gal4*) in vehicle fed flies: *ß-tub*>+ n=45*, ß-tub>Vha44-IR* n=56*, ß-tub>Vha68-2-IR* n=45*, ß-tub>Vha44* n=48*, ß-tub>VhaSFD* n=56. D) Enterocyte-specific depletion (*Myo1A-Gal4*) in vehicle fed flies. *Myo1A*>+ n=42, *Myo1A >Vha44-IR* n=58. Log-rank (Mantel–Cox) test; **** p<0.0001, *** p<0.001, ** p<0.01, n.s. p>0.05.

**S2 Fig. LAMP1-GFP is upregulated in the midgut in response to oxidative stress.** A-B') Representative images of LAMP1-GFP in R4-R5 midgut region of *Myo1A>UAS-LAMP1-GFP* adult males fed vehicle (A–A′) or 10 mM paraquat (B–B′). Scale bars: 25 µm. (C) Mean fluorescence intensity of LAMP1-GFP. Vehicle n=6, paraquat n=6. Student's t-test, ** p<0.01.

**S3 Fig. Vha44 depletion reduces enterocyte death following PQ exposure.** A-B′) Representative images of DCP-1 staining in the R4–R5 midgut region of Myo1A>+ (A, A′) and Myo1A>Vha44-IR (B, B′) flies fed 10 mM paraquat. Scale bars: 25 µm. C) Mean fluorescence intensity of DCP-1 staining. *Myo1A*>+ n=7, *Myo1A>Vha44-IR* n=6. Student’s t-test, ** p<0.01.

**S4 Fig. *Vha44*-dependent JNK activation is reproduced with an alternative oxidant.** A-B’) Representative images of pJNK staining in R4-R5 midgut region of *Myo1A*>+ (A, A′) and *Myo1A>Vha44-IR* (B, B′) flies fed 10 mM PQ. Scale bars: 25 µm. C) Mean fluorescence intensity of pJNK staining. *Myo1A*>+ n=5, *Myo1A>Vha44-IR* n=5. Student’s t-test, exact p-value shown. D-G) Representative images of *puc*-lacZ staining in the R4–R5 midgut region of vehicle (D, E) and 1% $H_2O_2$ (F-G) fed flies. Scale bars: 25 µm. H) Mean fluorescence intensity of *puc*-lacZ staining. Vehicle: *Myo1A*>+ n=11, *Myo1A>Vha44-IR* n=7; $H_2O_2$: *Myo1A*>+ n=9, *Myo1A>Vha44-IR* n=8. One-way Anova with Šídák's multiple comparisons test, *** p<0.001, n.s. p>0.05.

**S5 Fig. Paraquat induces *Relish*-GFP, and *Vha44* depletion suppresses paraquat-induced *Drs*-GFP.** A–B′) Representative images of the *Relish*-GFP reporter in the R4–R5 midgut region of *Myo1A*>+*;*

*Rel-GFP* flies fed vehicle (A–A′) or 10 mM paraquat (B–B′). Scale bars: 25 µm. C) Mean fluorescence intensity of *Relish*-GFP. Vehicle: *Myo1A>+; Rel-GFP* n=7; PQ: *Myo1A>+; Rel-GFP* n=7. Student's t-test, exact p-value shown. D–E′) Representative images of the *Drs*-GFP reporter in the R4–R5 midgut region of *Myo1A>+; Drs-GFP* (D, D′) and *Myo1A>Vha44-IR; Drs-GFP* (E, E′) flies fed 10 mM paraquat. Scale bars: 25 µm. F) Mean fluorescence intensity of *Drs*-GFP. *Myo1A*>+ n=4; *Myo1A>Vha44-IR* n=4. Student's t-test, ** p<0.01.
